# Further confirmation for unknown archaic ancestry in Andaman and South Asia

**DOI:** 10.1101/071175

**Authors:** Mayukh Mondal, Ferran Casals, Partha P. Majumder, Jaume Bertranpetit

## Abstract

In a recent paper^1^, we have derived three main conclusions: i) that all Asian and Pacific populations share a single origin and expansion out of Africa, contradicting an earlier proposal of two independent waves; ii) that populations from South and Southeast Asia harbor a small proportion of ancestry from an unknown extinct hominin – different from the Neanderthal and the Denisovan – which is absent in Europeans; and, iii) that the characteristic distinctive phenotypes (including very short stature) of Andamanese do not reflect an ancient African origin, but have resulted from strong natural selection on genes related to human body size. Although the single wave out of Africa^2^ and single origin for Asian and Pacific populations have been confirmed^3^, the existence of admixture with an extinct hominin has been challenged by Skoglund et al.^4^, as they were unable to replicate our results in their data sets. While we had used a wide variety of statistical methods and data sets from diverse populations to draw our inference, Skoglund et al.^4^ have used only one method (D-stats^5^, for the whole genome, not specifically for the relevant genomic regions) and compared only with the Asians, not even with the Europeans. Skoglund et al.^4^ have alleged that our statistical treatment of the data was faulty and have pointed out some possible sources of error. We have reexamined our data focusing on possible sources of error flagged by Skoglund et al^4^. We have also performed new analyses. The reexamination and new analyses have bolstered our confidence that our earlier inferences were correct and have resulted in an improved model of introgression of modern humans with a hitherto unknown archaic ancestry. We also propose a possible reason for the inability of Skoglund et al.^4^ to validate our inference.

---

In our reanalysis, first we test the impact on our inference of possible sources of bias, including the number of individuals, differences in coverage and batch effects. Second, we specifically address questions related to results in populations other than the Andamanese in relation to the detection of introgressed regions. In all cases we considered the ancestral position for humans as defined in 1000 Genomes Project as outgroup^6^ and not the chimpanzee, although the results are similar.

**1.- Number of Individuals.** Some of our analyses comprised 70 Indians compared to two Europeans, two East Asians and four Africans. This variation in sample size may have introduced a bias towards Indian-specific derived alleles. To check this, we have now performed variant calling using reads obtained by deep-sequencing of two Europeans, two Andamanese, one Papuan and two Africans together. Estimates obtained from Dstat analysis of these data are similar to those reported by us in Mondal et al^1^ and although the estimated values are slightly smaller, these values are similar to our original results and are still statistically significant (Table 1). Therefore, we rule out the possibility that because of heterogeneity in the number of individuals of different ethnicities included in our original analysis, we have made a false positive inference of introgression (at least for the Andamanese).

**Table 1:** Results of D-stats analysis using two individuals from each, Europe (French and Sardinian) and Africa (Yoruba and Mandenka), from an initial set of the SGDP^7^ and Andaman^1^ (Jarawa and Onge) and one Papuan^7^.

| W | X | Y | Z | D score | Z score |
| --- | --- | --- | --- | --- | --- |
| French | Jarawa | Yoruba | Ancestral | 0.0181 | 4.3 |
| French | Onge | Yoruba | Ancestral | 0.0158 | 3.427 |
| French | Jarawa | Mandenka | Ancestral | 0.0188 | 4.419 |
| French | Onge | Mandenka | Ancestral | 0.0135 | 2.936 |
| Sardinian | Jarawa | Yoruba | Ancestral | 0.0188 | 4.223 |
| Sardinian | Onge | Yoruba | Ancestral | 0.0166 | 3.543 |
| Sardinian | Jarawa | Mandenka | Ancestral | 0.0194 | 4.691 |
| Sardinian | Onge | Mandenka | Ancestral | 0.0142 | 3.166 |
| French | Papuan | Yoruba | Ancestral | 0.0336 | 6.893 |
| French | Papuan | Mandenka | Ancestral | 0.0269 | 5.439 |
| Sardinian | Papuan | Yoruba | Ancestral | 0.0344 | 6.847 |
| Sardinian | Papuan | Mandenka | Ancestral | 0.0277 | 5.782 |

**2.- Coverage.** Our data coverage is lower (15x) than in the comparison set (an initial set of the Simon Genome Diversity Project [SGDP]^7^). To test the possible impact of differences in sequencing coverage (as suggested by David Reich in a personal communication to us), we downgraded SGDP data to 15x coverage using -f flag in samtools^8^. The results (Table 2) are again statistically significant and are similar to those previously obtained^1^, suggesting that effect of differences in depth of coverage had not resulted in a false positive inference on introgression from an extinct hominin.

**Table 2:** Results of D-stats analysis using similar coverage for all individuals (15x). French, Sardinian, Yoruba and Mandenka are from an initial set of the SGDP^7^; Jarawa and Onge are from Mondal et al.^1^.

| W | X | Y | Z | D score | Z score |
| --- | --- | --- | --- | --- | --- |
| French | Jarawa | Yoruba | Ancestral | 0.015 | 3.945 |
| French | Onge | Yoruba | Ancestral | 0.0127 | 3.285 |
| French | Jarawa | Mandenka | Ancestral | 0.0129 | 3.202 |
| French | Onge | Mandenka | Ancestral | 0.0118 | 2.889 |
| Sardinian | Jarawa | Yoruba | Ancestral | 0.0167 | 4.011 |
| Sardinian | Onge | Yoruba | Ancestral | 0.0145 | 3.462 |
| Sardinian | Jarawa | Mandenka | Ancestral | 0.0144 | 3.677 |
| Sardinian | Onge | Mandenka | Ancestral | 0.0134 | 3.373 |

**3.- Batch effects due to laboratory of origin and processing of the sequences.** To check the impact of batch effects, we first downloaded the 1000 Genomes Project^9^ (KGP) vcf files and performed the D-stats analysis. The result obtained when we used Indian Telugu from the UK (ITU) to detect fewer shared African alleles using Yoruba in Ibadan, Nigeria (YRI) compared to Europeans using Utah Residents with Northern and Western Ancestry (CEU) [D(CEU,ITU;YRI,Ancestral)=−.0009 (Zscore=−0.933)] was not statistically significant. However, since data imputation was done in the KGP, we suspected that the statistically non-significant result that we obtained may be caused by having included imputed data. In fact, if Indian populations admixed with an unknown hominin population, the introgressed haplotypes would be at very low frequencies and therefore imputed results would be unreliable (a similar problem might arise if instead of sequence data, only SNP genotyping data would be used). We downloaded 10 CEU, 10 YRI and 10 ITU BAM files, converted to the same coverage (as the coverage is substantially divergent in this data set) and performed the variant calling without any imputation. The result we obtained was clearly in the same direction originally presented in Mondal et al^1^ and was statistically significant [D(CEU,ITU;YRI,Ancestral)=0.0079 (Zscore=3.747)]. Next, we used two CEU and two YRI (two individuals randomly chosen from KGP) with two Andamanese (AND) from our data, and also obtained a significant result (although the lower coverage of the KGP data markedly decreased the level of statistical significance) [D(CEU,AND;YRI,Ancestral)=0.0109 (Zscore=3.595)]. Then we retrieved fasta formatted data from the Simon Genome Diversity Project^3^ including one of each, Papuan, French, Sardinian, Dai, Han, Mandenka, Yoruba and Irula. We noted that variant-calling and filtering were already carried out on the raw data; we did not have the opportunity to carry these out ourselves. We obtained results similar to those presented in Mondal et al. (2016) for Papuan (Table 3), but highly variable and non-significant results for Dai, Han or Irula. While we have no way to identify the cause of these differences in results, we suspect that differences in treatment of raw data may be the contributing factors. We note that Mallick et al^3^ have used a new method for calling and filtering, which may have strong effects on calls of rare and interesting variants.

**Table 3:**
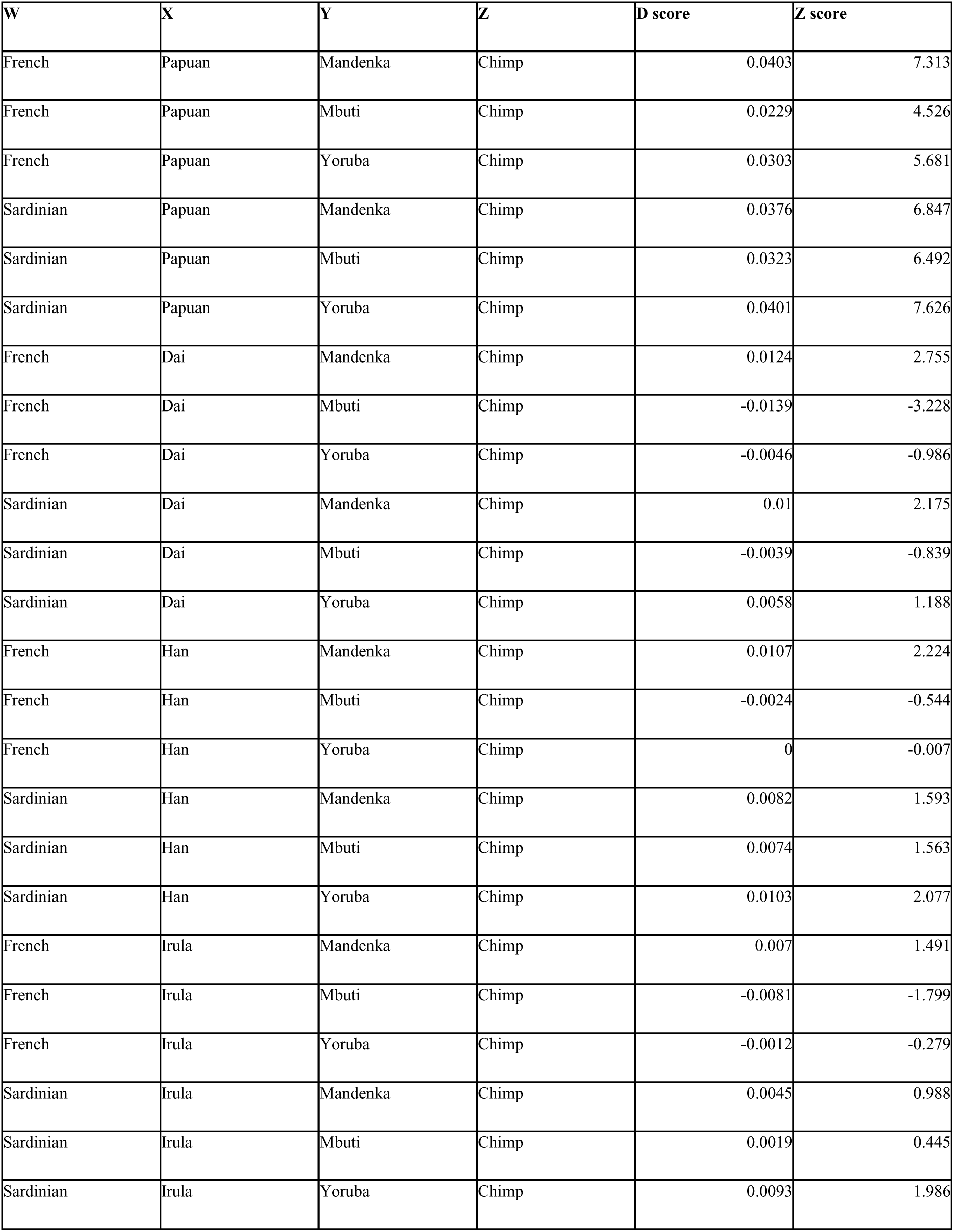
Results of D-stats analysis using data from SGDP (3rd data set)^7^: one Papuan, one French, one Sardinian, one Dai, one Han and one Irula.

| W | X | Y | Z | D score | Z score |
| --- | --- | --- | --- | --- | --- |
| French | Papuan | Mandenka | Chimp | 0.0403 | 7.313 |
| French | Papuan | Mbuti | Chimp | 0.0229 | 4.526 |
| French | Papuan | Yoruba | Chimp | 0.0303 | 5.681 |
| Sardinian | Papuan | Mandenka | Chimp | 0.0376 | 6.847 |
| Sardinian | Papuan | Mbuti | Chimp | 0.0323 | 6.492 |
| Sardinian | Papuan | Yoruba | Chimp | 0.0401 | 7.626 |
| French | Dai | Mandenka | Chimp | 0.0124 | 2.755 |
| French | Dai | Mbuti | Chimp | -0.0139 | -3.228 |
| French | Dai | Yoruba | Chimp | -0.0046 | -0.986 |
| Sardinian | Dai | Mandenka | Chimp | 0.01 | 2.175 |
| Sardinian | Dai | Mbuti | Chimp | -0.0039 | -0.839 |
| Sardinian | Dai | Yoruba | Chimp | 0.0058 | 1.188 |
| French | Han | Mandenka | Chimp | 0.0107 | 2.224 |
| French | Han | Mbuti | Chimp | -0.0024 | -0.544 |
| French | Han | Yoruba | Chimp | 0 | -0.007 |
| Sardinian | Han | Mandenka | Chimp | 0.0082 | 1.593 |
| Sardinian | Han | Mbuti | Chimp | 0.0074 | 1.563 |
| Sardinian | Han | Yoruba | Chimp | 0.0103 | 2.077 |
| French | Irula | Mandenka | Chimp | 0.007 | 1.491 |
| French | Irula | Mbuti | Chimp | -0.0081 | -1.799 |
| French | Irula | Yoruba | Chimp | -0.0012 | -0.279 |
| Sardinian | Irula | Mandenka | Chimp | 0.0045 | 0.988 |
| Sardinian | Irula | Mbuti | Chimp | 0.0019 | 0.445 |
| Sardinian | Irula | Yoruba | Chimp | 0.0093 | 1.986 |

**4.- Results in other Asian and Pacific populations.** Our study was mainly focused on the Andamanese population. We accept the suggestion that a more in depth introgression analysis should be performed in other populations using variant calls derived from a joint analysis of raw data. Regarding the Papuan and Australian populations, we think that the differences between them are related to variant calling artifacts, since only the vcf file was available for the Australian individuals. Further studies will have to consider the larger data set recently studied^2^. In relation to the East Asian populations, we did not get significant evidence of introgression, although the results of our analysis point to the possibility that East Asians also may have some introgression from this unknown hominin, even though the proportion of introgression may be lower than in the Andamanese.

**5.- Finding Introgressed Regions.** We have now performed analyses to detect introgressed regions in the Andamanese genomes using D-stats by genomic region and S* in our data. D-stats by region exhibited good statistical power to detect regions with strong divergence (that may have introgressed) from the simulated model (Figure 1) described in the methods section of Mondal et al^1^. It is clear that the differences in distributions of the D-stats by position correctly separated the two (introgressed and non-introgressed) distributions. If the results of D-stats and S*^10^ are combined, the extent of false positives decreased even further (Figure 2). The vertical line indicates the significance threshold for S*. The yellow dots in the graph mark the introgressed regions detected by the combined method, positive for both D-stats by position (defined as regions having −1 value) and S* (defined as regions having more than 95 percentile of the distribution of the non-introgressed model).

**Figure 1:**
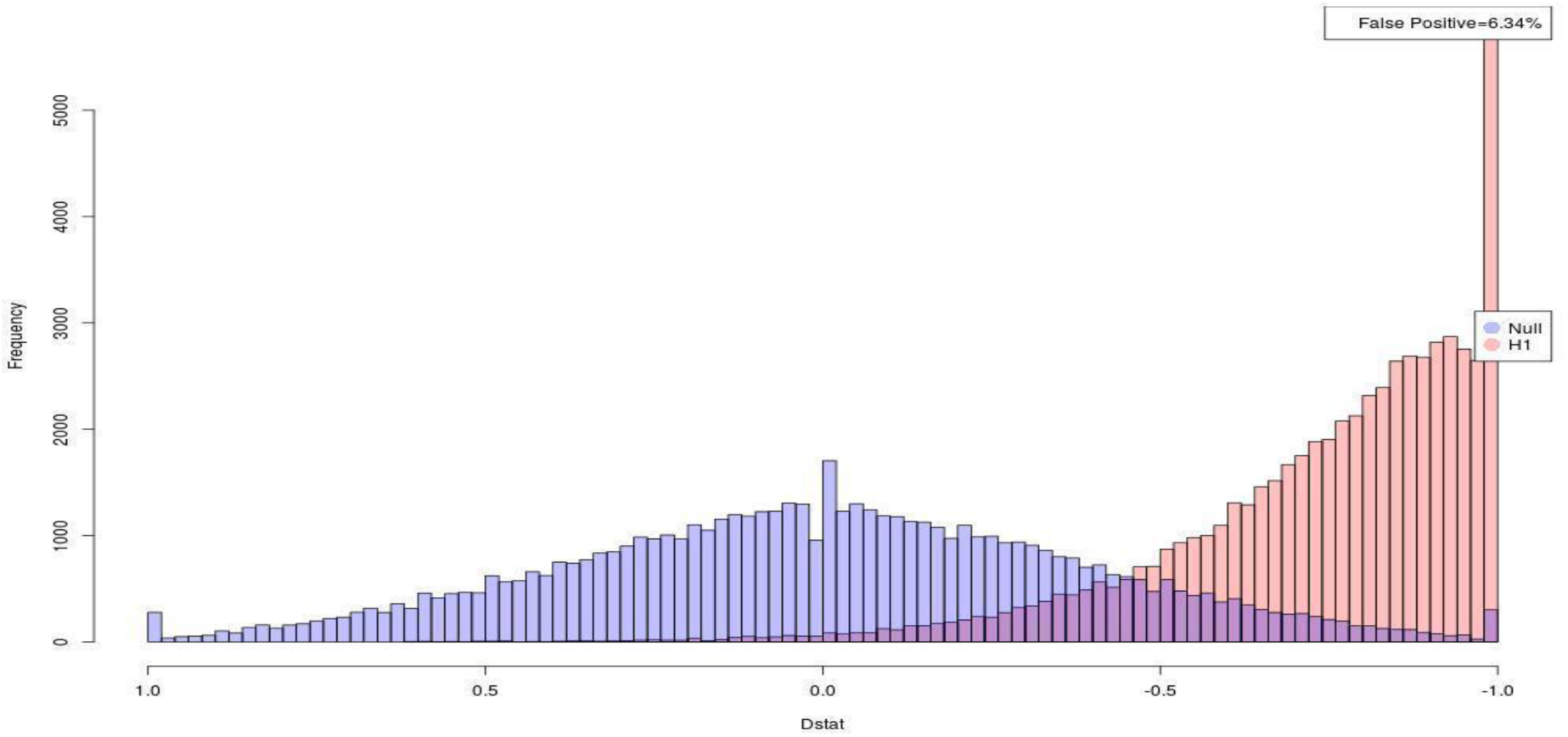
Efficiency of detecting putative introgressed regions by D-stats. The blue bars represent the standard distribution without introgression, and the red bars represent the divergent distribution that may have introgressed; both cases refer to a 50Kb region simulated 60,000 times. Not drawn to scale, as the red distribution would be much less frequent than the blue one in a scenario of small amount of introgression.

**Figure 2:**
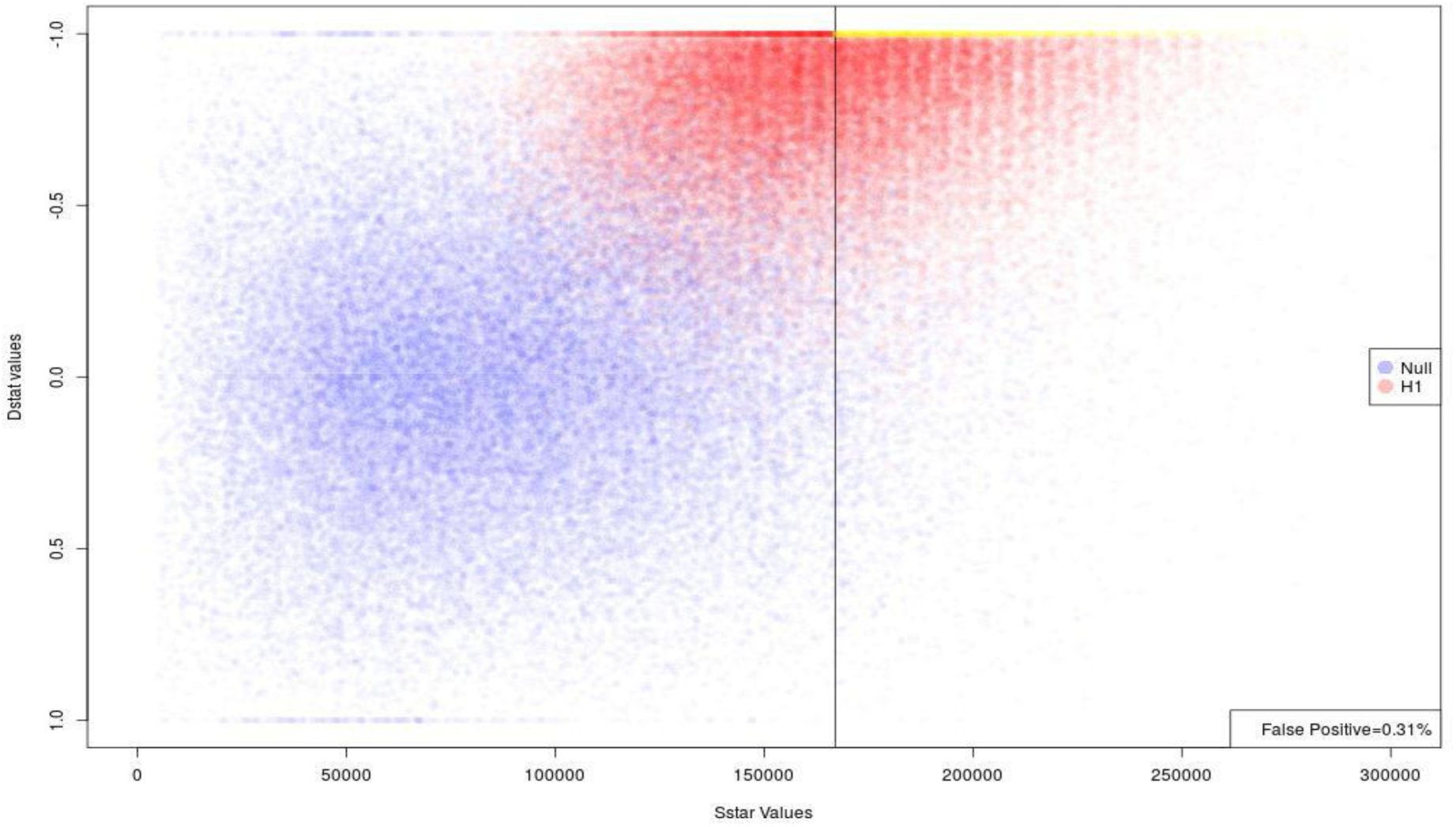
Efficiency of detecting putative introgressed regions by D-stats and S* in the simulations as in Figure 1. The blue points are the non-introgressed regions, the red points are the regions with high divergence, and yellow points are the regions that are extreme for both D-stats and S* and are thus considered introgressed.

Finally, we analyzed the regions which were more likely to be introgressed for both D-stats and S* (which accounted for a total of ∼15Mb per individual), by comparing to other hominin sequences and resulted clearly in a sequence basal to the Neanderthal-Denisovan clade (Figure 3).

**Figure 3:**
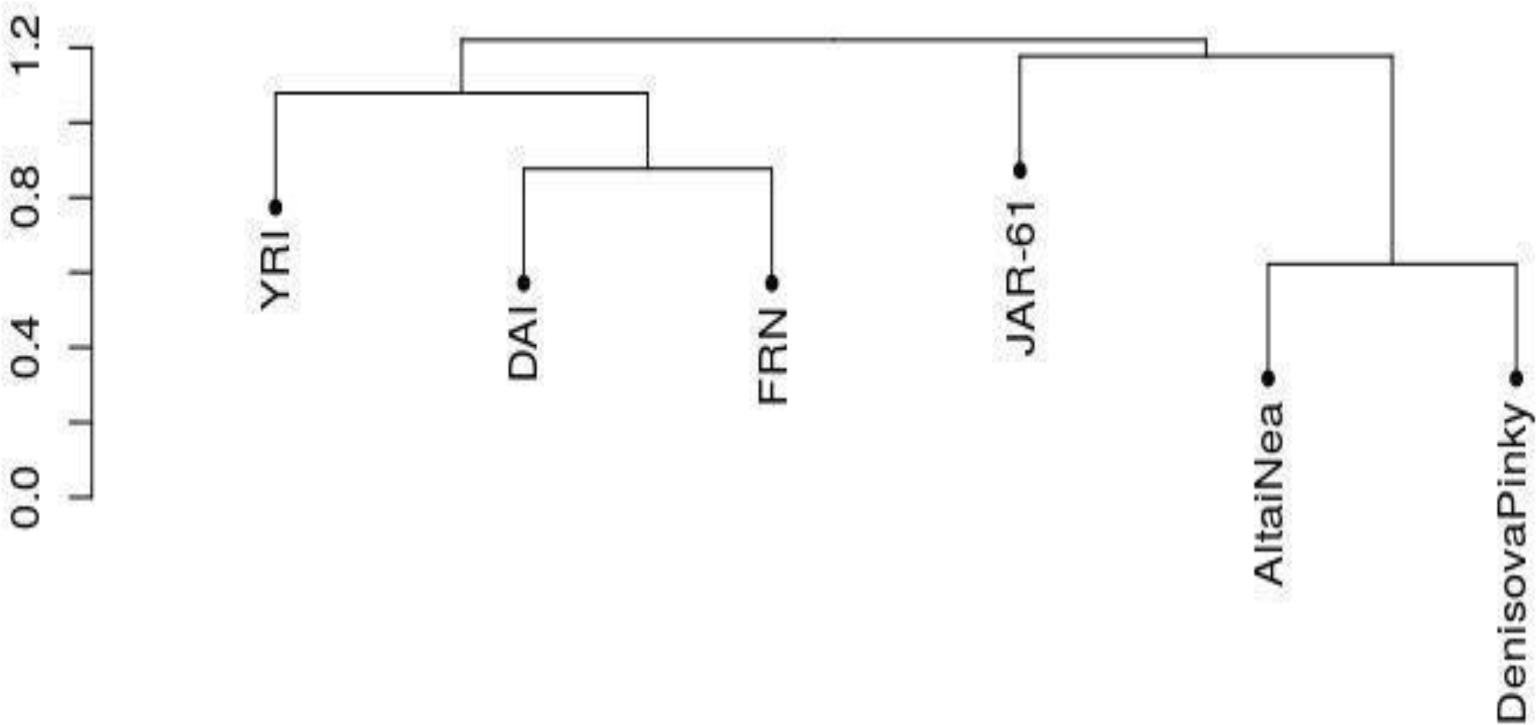
Hierarchical cluster analysis on the dissimilarity matrix of individual sequences. Modern humans: YRI, Yoruba, DAI from China, French, French. Other sequences: JAR-61, putative introgressed regions in a Jarawa, AltaiNea, Neanderthal from Altai, DenisovaPinky, Denisova.

## Discussion

To the extent feasible, we have examined all putative sources of bias^4^ – such as batch effects, sample size and coverage differences, data processing artifacts – that may have impacted on our earlier inference^1^ of the presence of an unknown hominin ancestry in the Andamanese population. We have shown that our earlier inference was robust and that we had not made a false-positive inference. We have replicated the results originally presented by us^1^ using additional data, independently of the origin of the data, when we were able to obtain raw data and analyze the data using a common set of methods. We wonder whether the new method of calling and filtering raw data introduced by Mallick et al^3^ and likely used by Skoglund et al^4^ may be the reason for the differences in results obtained by Skoglund et al^4^ and Mondal et al^1^. In addition, results of the present analyses of the introgressed regions further support our earlier conclusions^1^.

Finally, since in population genetics studies, disparate data sets generated by various laboratories are used, we propose that, in a manner similar to what has been done for somatic mutation detection in cancer^11^, the population genetics research community should initiate a joint effort to study the impact of using different variant-calling and filtering procedures on population genetic inferences. We are, of course, willing to contribute data and computing effort to such an endeavor (which could comprise a subset of the sequences of the SGDP^3^).

## Methods

### Variant Calling

The BAM files used were previously generated^1^ or downloaded from 1000 Genomes Project site^9^. In some of the analysis we downgraded the coverage of BAM files using -f flag from samtools-1.2^8^. Variant calling was done by GATK-3.5^12^ closely following “best practices of GATK”. Calling was done with default options and using --max-alternate_alleles 20 (to capture all genetic diversity present in the populations) by running HaplotypeCaller in GVCF mode on each sample separately. Then GenotypeGVCFs was used to produce the joint genotyping of the gVCFs on all of the samples together to generate a raw SNP and indel Variant Calling File (VCF). The raw VCF was filtered using post variant calling recalibration steps as listed in GATK “Best Practices”. VariantRecalibration from GATK were used to calculate various statistics for novel variants (both for SNPs and indels) and then recalibrated according to their needs using ApplyRecalibration. As different data sets have differing amounts of false positives, it was necessary to give the appropriate importance to the different data sets for the training algorithm. We took the value for prior likelihood of true sites defined in the GATK website for the corresponding data sets.

For the recalibration steps, the following datasets were used, with the following commands: SNPs

1. dbsnp version 137: -resource:dbsnp, known=true, training=false, truth=false, prior=2.0.
2. hapmap version 3.3: -resource:hapmap, known=false, training=true, truth=true, prior=15.0 (International Hapmap3 Consortium 2010).
3. Omni genotyping array 2.5 million 1000G: -resource:omni, known=false, training=true, truth=true, prior=12.0.
4. 1000G phase 1 high confidence: -resource:1000G, known=false, training=true, truth=false, prior=10.0.
5. With the flags -an QD -an MQRankSum -anReadPosRankSum -an FS -an DP -an InbreedingCoeff. All other parameters were set to default values.

Indels

1. Mills 1000G high confidence indels: -resource:mills, known=false, training=true, truth=true, prior=12.0.
2. dbSNP version 137: -resource:dbsnp, known=true, training=false, truth=false, prior=2.0.

With the flags --maxGaussians 4 -an FS -an ReadPosRankSum -an MQRankSum -an DP -an InbreedingCoeff. All other parameters were set to default values.

### D-stats Calculation

We first converted the vcf file to plink format using vcftools^13^ and we added ancestral information, which was downloaded from 1000 Genomes Project site, using plink^14^. We converted plink to eigensoft format using convertf and D-stats was calculated using qpDstat both of them came together with Admixtools 1.1^5^. All the D-stats calculations were done on SNPs which are present in every Individual thus not biasing for allele frequency which present in low covered regions.

### SGDP 3^rd^ dataset

We downloaded SGDP files from the site http://sharehost.hms.harvard.edu/genetics/reich_lab/cteam_lite_public3.tar, 11/07/2016 Individual information was extracted using cpoly^3^. Individuals used in this analysis were: Chimp, B_French-3, B_Mandenka-3, B_Mbuti-4, B_Yoruba-3, S_Irula-1, B_Papuan-15, B_Sardinian-3, B_Dai-4 and B_Han-3.

### Simulations

We used ms to simulate demographic models^15^. The Andamanese demographic model is based on Mondal et al^1^.

To calculate efficiency, two different models were used. 1) The null model: we used Andamanese demography with modern humans without using any introgression from hominin. 2) The H1 model: we simulated a hominin population which has diverged from modern humans 300 kya ago (as mutation rate 2.3×10^−8^ was used). This is the best scenario. In both cases we simulated around 50 kb of 60,000 regions. We chose a specific number of segregating sites of 100 and recombination rate of 1.3×10^−8^.

Null=ms 20 60000 -s 100 -r 26 50000 -I 3 4 4 2 -n 1 1.4474 -n 2 3.3814 -n 3 .1055 -m 1 2 1.244 -m 2 1 1.244 -m 2 3 0.88 -m 3 2 0.88 -g 2 179 -ej .023 3 2 -en .023 2 .1861 -em .023 1 2 6 -em .023 2 1 6 -em .023 1 3 0 -em .023 3 1 0 -em .023 2 3 0 -em .023 3 2 0 -ej .051 2 1 -en .148 1 .73 -p 5

H1=ms 10 60000 -s 100 -r 26 50000 -I 3 4 4 2 -n 1 1.4474 -n 2 3.3814 -m 1 2 1.244 -m 2 1 1.244 -g 2 179 -en .023 2 .1861 -em .023 1 2 6 -em .023 2 1 6 -em .023 1 3 0 -em .023 3 1 0 -em .023 2 3 0 -em .023 3 2 0 -ej .051 2 1 -en .148 1 .73 -ej .4 3 1 -p 5

D-stats^5^ and S*^10^ was calculated as mentioned in their respective.

### Data repository

vcf files from Andamanese, Indian and a set of comparison from an initial set of the SGDP^7^ are available temporarily in: https://www.dropbox.com/s/rk6jjjocefhwns1/GreatApe_Indiafinal_Austosome.vcf.gz?dl=0 and will be uploaded soon in a general server.

